# Predicting the Sound-Induced Flash Illusion: A Time-Window-of-Integration Approach

**DOI:** 10.1101/2025.03.19.644110

**Authors:** Hans Colonius, Adele Diederich

**Affiliations:** Department of Psychology, Carl von Ossietzky Universität, Oldenburg, Germany and Department of Psychological Sciences, Purdue University; Department of Medical Physics, Carl von Ossietzky Universität, Oldenburg, Germany and Department of Psychological Sciences, Purdue University

## Abstract

The sound-induced flash illusion (SIFI) refers to the observation that pairing a single flash with 2 auditory beeps leads to the illusory perception of 2 visual stimuli (fission illusion). Susceptibility to the illusion depends on many factors like exact physical stimulus parameters, participants’ expectations, attention, age, and clinical/non-clinical population membership. While there exist many experimental studies, very few models have been proposed to account for the phenomenon. Here we suggest a formal model (SIFI-TWIN) based on the notion of a temporal binding window that predicts the occurrence of illusory flashes as a function of the temporal and physical stimulus arrangement. The model’s performance is illustrated on a study investigating differences in SIFI performance between elderly hearing aid users and those with the same degree of mild hearing loss who were not using hearing aids. The results suggests that the higher incidence of reporting an illusory flash for hearing-aid users is due to both a larger temporal window of integration and a larger bias to report the illusion. The SIFI-TWIN model will help to better understand the diverse results from clinical and non-clinical studies as well as the cognitive foundations of the SIFI.

---

Integrating information from multiple senses typically improves performance in many different perceptual tasks including target detection or discrimination (Stein & Meredith, 1993; Diederich & Colonius, 2004; Drugowitsch, DeAngelis, Klier, Angelaki, & Pouget, 2014). The effect holds for some higher-level cognitive tasks as well. For example, it has been known since long (Erber, 1969) that lipreading improves speech understanding: most adults benefit from being able to see the talker’ s face when speech is degraded (for a recent review, see Bernstein, Jordan, Auer, & Eberhardt, 2022). On the other hand, presenting conflicting stimuli across modalities often results in deceiving the observer; well-known examples are the ventriloquist effect (Bertelson & Radeau, 1981) and the McGurk-McDonald effect (McGurk & McDonald, 1976). Another case in point is the popular *sound-induced flash illusion* (SIFI).

In its simplest version, SIFI refers to the observation that pairing a single flash with 2 auditory beeps leads to the illusory perception of 2 visual stimuli, referred to as *fission illusion* (see Figure 1). It has been demonstrated that susceptibility to the illusion depends on many factors, like exact physical stimulus parameters, participants’ expectations, attention, age, and clinical/non-clinical population membership; for a recent review, see Hirst, McGovern, Setti, Shams, and Newell (2020). Since the paper by Shams, Kamitani, and Shimojo (2000), the last 25 years have seen a huge amount of theoretical and empirical results collected on the effect. While there are many experimental studies, very few models have been proposed to account for the phenomenon. Here we suggest a simple model based on the notion of a temporal binding window.

**Figure 1.**
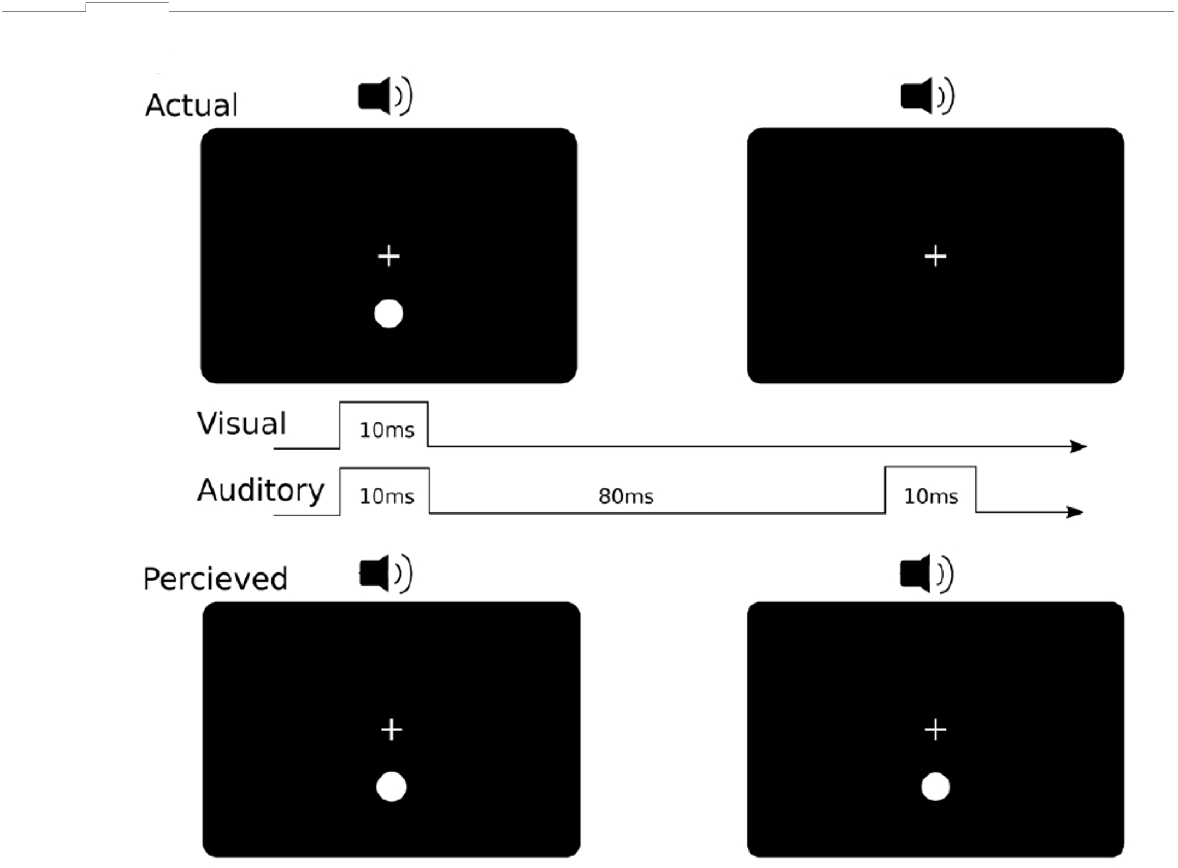
Example illusory condition in the SIFI paradigm in which one flash is presented with two beeps (upper), typically producing the perception of two flashes(lower). Visual stimuli are typically presented for the duration of 1 frame (10 ms on a 100 Hz monitor) (after Figure 1 from (Hirst et al., 2020))

In a generalization of the original SIFI setup, Wozny, Beierholm, and Shams (2008) presented multiple (e.g., 0 to 2) visual, tactile, and auditory stimuli in various combinations and subjects were required to report the number and type of stimuli of presented. Similarly, Odegaard, Wozny, and Shams (2016) varied the number of flashes and beeps from 1 to 4. Their results were best described in terms of the optimal Bayesian causal inference (BIC) model (Rohe, Hesse, Ehlis, & Noppeney, 2024; Körding et al., 2007; Shams & Beierholm, 2022). Specifically, the final percept (how many beeps and/or flashes?) is based on (i) the prior probability of a common cause for both beeps and flashes, (ii) the expected number of stimuli presented, and (iii) reliability of the sensory signal.

Given the close temporal proximity of cross-modal stimuli presented in a given trial, SIFI is commonly considered as an indicator of the strength of multisensory integration (MI). Importantly, one major factor determining the probability of observing a fission response is the time interval between the occurrence of the first beep (together with the flash) and the second beep; here this interval is referred to as *stimulus onset asynchrony* (SOA). Multiple studies from a variety of populations (see Figure 2) differing in age, clinical condition, and training, consistently showed that the probability of a fission illusion peaks at SOAs of about 70 to 100 ms and then declines in a nearly monotonic manner from values of around 70-80 % down to 10% (e.g., Chan, Connolly, & Setti, 2018; Bidelman, 2016).

**Figure 2.**
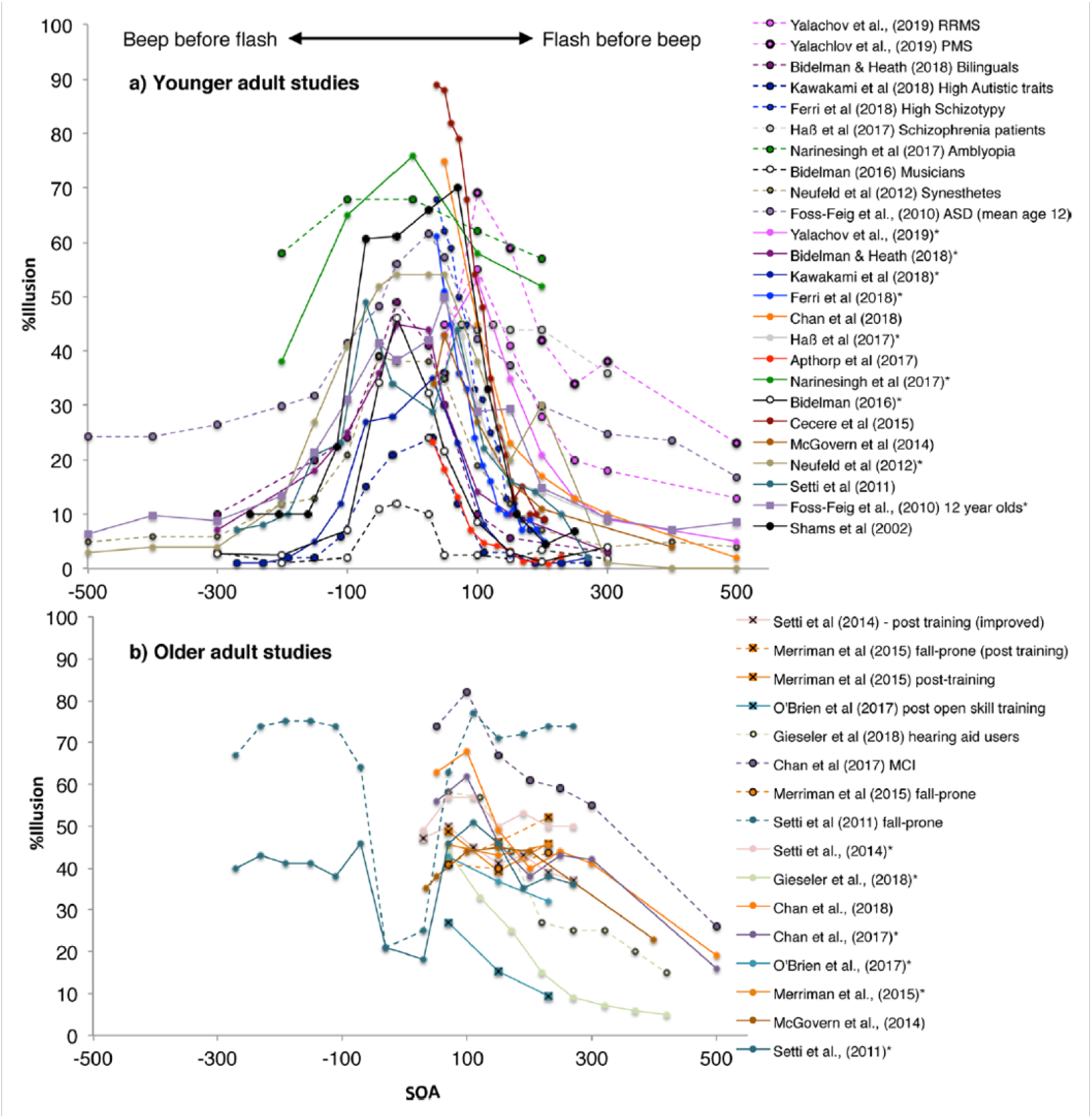
Studies using Stimulus-Onset Asynchrony(SOA) in ms used to assess SIFI susceptibility (after Figure 4 from (Hirst et al., 2020))

While this observation is in accordance with the BIC explanation, the underlying mechanism is not made explicit in the model. To fill this gap, here we suggest a simple mechanism that predicts the probability of an illusion to occur as a function of the temporal and physical stimulus arrangement, called time-window-of-integration model for the sound-induced-flash-illusion, or SIFI-TWIN model, for short. It is based on the popular “temporal binding window” concept according to which auditory and visual stimuli are perceived as integrated events whenever they co-occur within this window (see, e.g., Stevenson, Zemtsov, & Wallace, 2012; Colonius & Diederich, 2004).

We first introduce our basic assumptions about the time window mechanism underlying the fission illusion in an informal way. Then we present the model in more detail and derive specific predictions for a parametric version of the model. Finally, the model’s performance is illustrated on a data set from our lab comparing the strength of the illusion between users and non-users of a hearing-aid.

## TWIN–SIFI model

(1) *Each stimulus triggers a unique internal process unfolding over time that can be represented by a random variable*. Representing the processing time triggered by a stimulus presentation by a random variable is common in cognitive modeling; here it may be interpreted as peripheral, neurosensory processing time without specifying the associated distribution function.
(2) *The two auditory stimuli (beeps) are registered in correct temporal order when the time between them is large enough*. The individual minimum time required can be estimated, e.g., by presenting 2 successive beeps from different directions.
(3) *After presentation of the first beep and the flash, their processing times “race” against each other; the “winner” opens a “time window of integration” of a specific length*. The concept of “opening the time window” simply means that the temporal distance to the second beep, i.e. the SOA, is measured from the time of registration of the winner of the race, either the flash or the first beep. No claim is made about the level of awareness involved in “registering” the winner and termination order of two processes.
(4) *There are three mutually exclusive conditions for integration: (i) the flash opens the window and processing of both beeps terminates within the window; or, (ii) the first beep opens the window and processing of the flash terminates before the second beep within the window or, (iii) processing of the second beep terminates before the flash within the window*. This means that, if temporal proximity is satisfied, visual and auditory information are integrated into a multisensory event so that the distinction between the unisensory percepts is no longer available.
(5) *Participant reports seeing an (illusory) second flash if (i) one of the events in Assumption (4) occurs or, (ii) if no integration occurs but it is nevertheless reported (response bias)*. An illusory flash is reported not only when multisensory integration occurs but also if participant has a tendency to report it, e.g., because of a high prior probability of a second flash being presented in the experimental context.

### Formal Model Specification

The basic assumptions are specified as follows; their numbering corresponds to the one above.

(S1) *There exist jointly distributed non-negative, continuous, and stochastically independent random variables V, A*_1_, *and A*_2_ *denoting the processing time of the flash, the first and second beep, respectively*. Assuming stochastic independence among all random variables seems plausible as a first step in order to keep derivations simple but it can, in principle, be dropped by adding some stochastic dependencies.
(S2) *With τ* (*τ ≥ τ*_0_ *>* 0) *denoting the SOA value, P* (*A*_1_ *< A*_2_ + *τ*) = 1. Choosing the value of *τ*_0_ always large enough^1^ guarantees that *P* (*A*_1_ *< A*_2_ + *τ*) = 1.
(S3) *P* (*A*_1_ *< V*) + *P* (*V < A*_1_) = 1. This simply states that either the first beep or the flash opens the window.
(S4) *Integration happens if, and only if, one of the following 3 (mutually disjoint) events occurs:*

(i) *E*_1_(*τ, ω*) ≡ *{A*_1_ *< V < A*_2_ + *τ < A*_1_ + *ω}, or*
(ii) *E*_2_(*τ, ω*) ≡ *{A*_1_ *< A*_2_ + *τ < V < A*_1_ + *ω}, or*
(iii) *E*_3_(*τ, ω*) ≡ *{V < A*_1_ *< A*_2_ + *τ < V* + *ω}* *with ω denoting the window width. We will assume throughout that ω >* 0. These are the only 3 stochastic events under which (sufficient) temporal proximity among the termination times of the flash and the beeps leads to integration; note that window width parameter *ω* has to be estimated from the data, whereas *τ* is specified by the experimental setup. We denote the probability of the union of the three events as *p*_*I*_(*τ* ; *ω*):

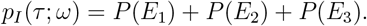

where we dropped the dependency of *E*_*i*_ (*i* = 1, 2, 3) on *τ* and *ω* for simplicity. (S5) *The probability that participant reports the illusion when SOA equals τ is*

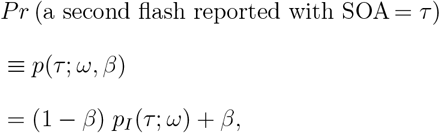

*where β* (0 ≤ *β* ≤ 1) *is a bias parameter denoting the probability to respond seeing a second flash when no integration has occurred and p*_*I*_(*τ* ; *ω*), as above, *the probability of integration*. For *β* → 0 the response approximates the probability of integration *p*_*I*_(*τ* ; *ω*), whereas for *β* → 1 the response is increasingly determined only by a bias.^2^

Figure 3 depicts a schematic of the time window mechanism.

**Figure 3.**
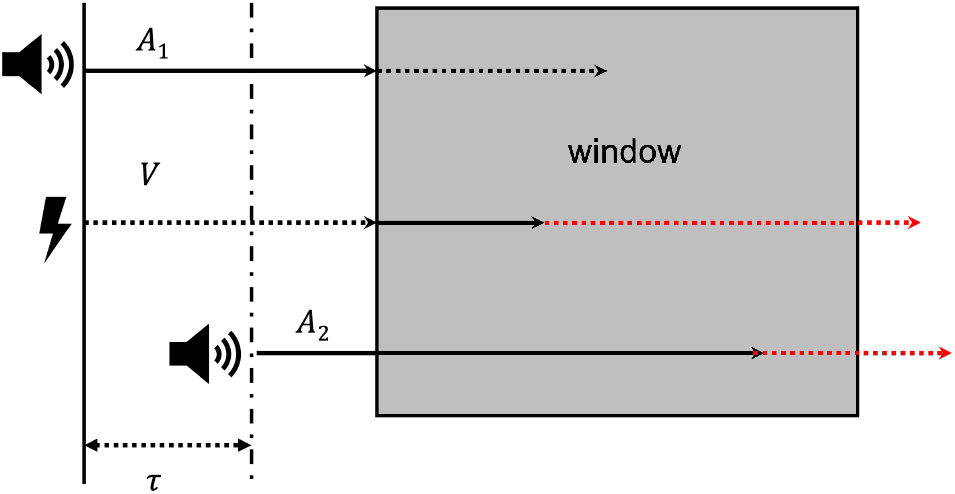
A schematic of the time window mechanism; A_1_, A_2_ refer to beep processing times, V to flash processing time. An illusion occurs if and only if all times terminate within the window.

### Computing the probability of reporting an illusory flash (fission illusory)

In order to predict the probability of an illusion, we have to compute the probability of each of the 3 events mentioned in (S5). Assumption (S1) implies that there exists a probability density function (pdf) for each of the random variables involved in the event: denote the pdf of *A*_1_ by *f*_1_(*x*_1_), the pdf of *A*_2_ *f*_2_(*x*_2_), and the pdf of *V* by *f*_3_(*x*_3_). Starting with the first event, we get

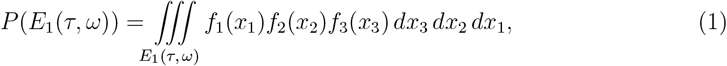

where integration is across the subset of positive real points (*x*_1_, *x*_2_, *x*_3_) in 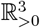 (ℝ_*>*0_ the set of positive real numbers) defined by

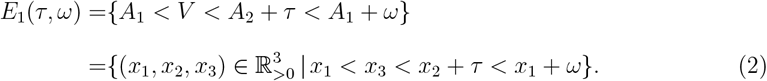

Expressions for the other 2 events are similar. For obtaining a quantitative prediction of the probability of integration, we need to specify the 3 pdfs. Keeping computations simple, the first choice is the exponential distribution, so that for *i* = 1, 2, 3,

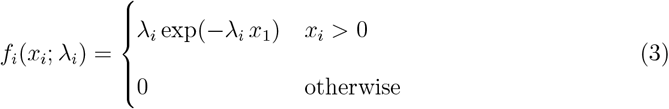

with parameters *λ*_*i*_ *>* 0. Evaluating the integrals under this assumption is elementary; nonetheless, their sum adds up to a lengthy expression for the probability of integration *p*_*I*_(*τ* ; *ω*; *λ*_1_; *λ*_2_; *λ*_3_) as function of the parameters *ω, λ*_1_, *λ*_2_ and *λ*_3_ (given in the appendix). The probability of integration greatly simplifies by setting all distribution parameters equal: with *λ*_1_ = *λ*_2_ = *λ*_3_ ≡ *λ* we get

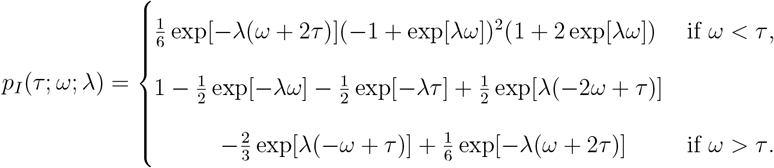

For this case, Figure 4 depicts integration probability as a function of SOA (*τ*) and window width (*ω*). Delaying presentation of the second beep (i.e., *τ* increasing from 70 to 420 ms), in general decreases the probability of integration towards zero. This is consistent with the notion that the window no longer captures a second beep that comes in too late^3^.

**Figure 4.**
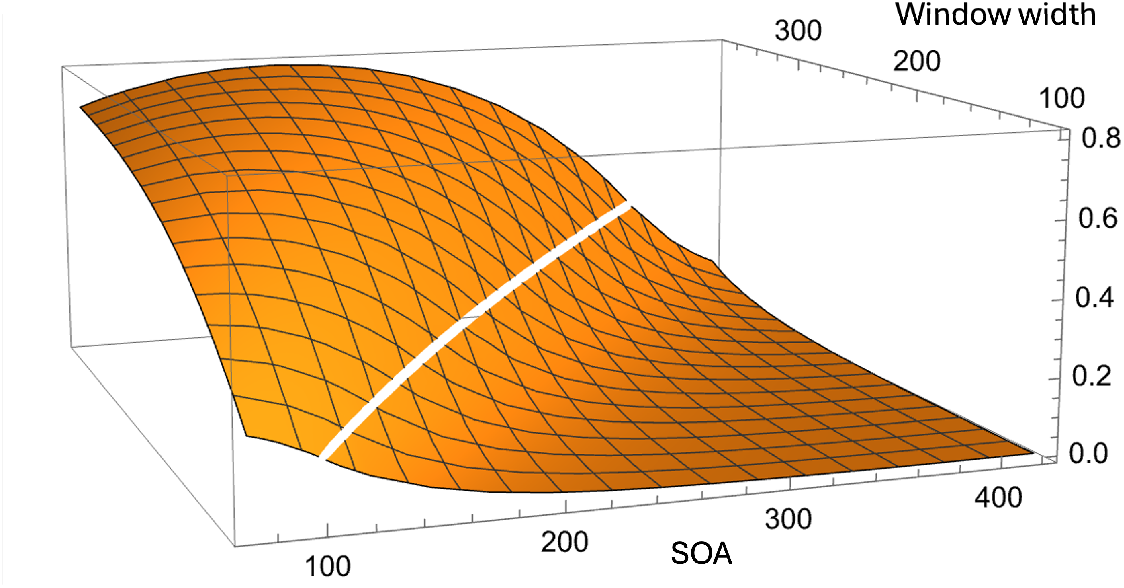
Probability of integration *p*_*I*_(*τ* ; *ω*; *λ*) with exponential pdfs as a function of SOA = *τ* ∈ (70, 420) and window width = *ω* ∈ (100, 370) with common density parameter *λ* set to *λ* = .01 in the TWIN–SIFI model.

Finally, a participant with bias parameter *β* reports an illusory flash with probability

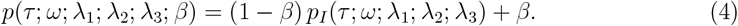

Whether assuming equality of the distribution parameters is plausible depends on the experimental setup and may have to be dropped. In particular, some studies investigate the effect of varying the intensity of the second beep relative to the first one suggesting different lambda values for the beeps (e.g., Ito, Matsumoto, & Kobayasi, 2023).

### Effect of hearing-aid use on SIFI

Gieseler, Tahden, Thiel, and Colonius (2018) were interested in the possible effect of devices for hearing support, such as hearing aids, on multisensory processing.^4^ They investigated differences in SIFI performance between elderly hearing aid users (HU) and those with the same degree of mild hearing loss who were not using hearing aids, the non-users (NU). It is well-known that the auditory modality affords superior temporal discrimination, and we assume that this would persist in age-related mild hearing loss. Thus, both groups should place more weight to audition relative to vision, which would render both susceptible to the flash illusion induced by the presence of the beeps. Moreover, the NU group experiencing degraded auditory input in everyday life might put less weight on audition and, thus, should show less integration than the HU group which had their hearing abilities restored (but were tested without hearing aids).

Comparing two groups of equal sample size (*n* = 33) across 8 different SOAs, Gieseler et al. (2018) did in fact find a higher incidence of the illusion in the HU group, even when various control conditions were taken into account (for more details including signal detection analyses, the original paper needs to be consulted). Both panels in Figure 5, depicting all data across subjects, show that the illusion for group HU was consistently above that of group NU across all 8 SOA value.s

**Figure 5.**
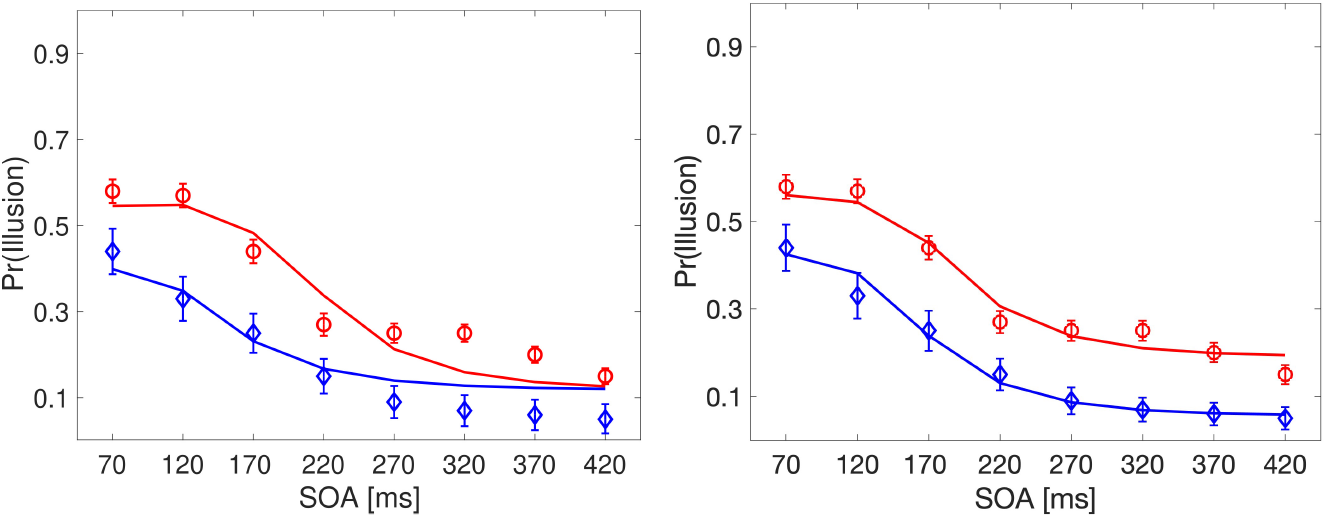
Predicted probability (lines) and observed relative frequency (symbols with 95% confidence intervals) of reporting a second flash (fission illusion). HU group values (open circles) are consistently above those of NU group (open diamonds). *Left panel*: fit of TWIN–SIFI model *λ*-*ω*_*HU/NU*_ -*β* allowing different integration windows: *ω*_*HU*_ = 218, *ω*_*NU*_ = 145, and *β* = .12. *Right panel*: fit of TWIN–SIFI model *λ*-*ω*_*HU/NU*_ -*β*_*HU/NU*_ allowing both different integration windows and different bias parameters: *ω*_*HU*_ = 189, *ω*_*NU*_ = 162, *β*_*HU*_ = .19 and *β*_*NU*_ = .06; see Appendix B for model notation.

Next, we probed the TWIN–SIFI model with exponential distributions on their data. Parameters were estimated minimizing the weighted sum of squares between predicted and observed frequencies. Five model versions with different sets of parameters were fitted and compared via the root mean square error of approximation (RMSEA) that takes the number of parameters into account. While none of the models reached a satisfactory fit (e.g., Schubert, Hagemann, Voss, & Bergmann, 2017), the model version with separate parameters for window size (*ω*) and bias (*β*) fared best (for details, see Appendix B).

The *left panel* shows best-fitting curves monotonically decreasing with SOA for windows size *ω*_*HU*_ = 218 and *ω*_*NU*_ = 146 that reflect the higher integration probability of HU compared to NU. Stimulus parameter *λ* = .009 and bias parameter *β* = .12 were estimated at the same time but held equal for both groups.

For SOA values larger than 270, there is a tendency to underestimate the HU values and to overestimate the NU values, as shown in the left panel. However, when the bias parameter for both groups was allowed to differ during the estimation procedure, these tendencies disappear, with *β*_*HU*_ = .19 and *β*_*NU*_ = .06 (see right panel of Figure 5). This is in line with the notion that for SOA values of 270 ms or larger, the probability of integration should be relatively small given the window size and, thus, participants’ illusion responses should increasingly be determined by bias. Moreover, the improvement obtained by using separate bias parameter estimates –the value for HU is about three times larger than the one for NU– suggests that hearing-aid users have a higher prior probability for assuming a common cause of beeps and flashes, in accordance with the Bayesian causal inference model. This is arguably a consequence of hearing-aid users more often experiencing the stimuli as a common visual-auditory event, due to their improved acoustic faculty, but this speculation requires further experimental back-up.

## Discussion

The sound-induced flash illusion (SIFI) is increasingly recognized as an important tool for measuring multisensory integration. In their review, Hirst et al. (2020) pointed out that the most critical parameter modulating the SIFI may be the time between the first beep/flash pair (stimulus onset asynchrony, SOA). Extending the Bayesian causal inference explanation, the TWIN–SIFI model introduces a mechanism specifying the effects of SOA on the illusion. It is based on the notion of a temporal window determining whether or not an integration of visual and auditory events –flashes and beeps– occurs in a given trial. Introducing a race mechanism, the probability of integration for a given SOA becomes a function of the individual intensity parameters of beeps and flashes. Finally, the probability of reporting a second, illusory flash is accounted for by combining a bias parameter with the integration probability. Up to now, the width of the temporal window has been defined in a more or less *ad-hoc* manner, for example, as “the contiguous span of successive SOAs in the 1 flash–2 beeps condition in which the proportion of illusory reports is significantly greater than this proportion in the 1 flash–1 beep control condition” (Foss-Feig et al., 2010). In contrast, the TWIN–SIFI model features window width as a built-in parameter, *ω*. In a similar vein, the bias to report an illusory flash is usually defined relative to the frequency of reporting 2 flashes in the 1 flash–1 beep control condition, whereas here bias is uniquely defined as another model parameter, *β*.

The model can be tested by experimentally by varying any factor that should affect these parameters. We illustrated this with a study probing the effect of hearing-aid use on the SIFI. This analysis suggested that the higher incidence of reporting an illusory flash for hearing-aid users is due to both a larger temporal window of integration and a larger bias. The multitude of studies where both the SOA and some other factor are varied (see Figure 2) provides a convenient testing ground for further tuning the model. We believe that this will help to better understand the often mixed results from clinical and non-clinical studies as well as the cognitive foundations of the SIFI.

Obviously, the current version of the TWIN–SIFI model has a number of limitations. Assuming exponential distributions has only been justified by the ease of deriving predictions and may have to be modified if suggested by data. Given that in studies with younger adults, the beep is often presented at various time points before the flash (negative SOAs, see upper panel of Figure 2), it is necessary to modify the model equations to comprise negative SOAs as well (e.g., by reversing the role of the first and second beep). Moreover, the model should be extended to cover situations where more than 2 beeps and flashes are being presented within a trial (e.g., Odegaard et al., 2016).

There are several other experimental setups the TWIN–SIFI model could possibly be adapted to. One, seemingly closely related to the SIFI, is the *fusion illusion*, that is, reporting one flash when two flashes are presented with one beep. However, such fusion effects are less reliable compared with fission effects, and they have been studied far less than the former; for further discussion of the substantial differences between the effects, see Hirst et al. (2020).

A different, and more straightforward, extension of our model is an effect reported in Stiles, Li, Levitan, Kamitani, and Shimojo (2018). In the “illusory audiovisual rabbit” effect, the perceived location of an illusory flash is influenced by an auditory beep-flash pair that follows the perceived illusory flash. Specifically, with a sequence of [beep–flash, beep, beep–flash], an illusory flash is perceived during the second beep. While all beeps are presented in the same central location, the illusory flash is perceived mid–way between the first and last flash, which are 2.84 degrees apart. Our preliminary analysis, implementing this setup in a schematic analogous to the one in Figure 3, indicates that 10 different stochastic outcomes of the race, instead of the 3 stochastic events for the SIFI, would have to be dealt with in the model.^5^

Because in the “audiovisual rabbit” the illusory flash location is determined by the location of the beep–flash pair presented *after* it, the authors consider this effect as an example of “perceptual postdiction”. In their words, *“Perceptual postdiction provides a relatively straightforward solution to this seeming paradox, as conscious awareness progresses neither simultaneously nor synchronously with the presentation of sensory stimuli. Instead, awareness is delayed in time to allow for the integration of sensory processing from earlier stimuli. Consequently, the brain can process a series of sensory inputs as a group before an individual becomes aware of them*.*”* (Stiles, Tanguay Jr., & Shimojo, 2021). Note that this dovetails neatly with our notion that “registering” the winner of the race and determining the order of the processes within the temporal window does not necessarily require awareness.

This raises the issue of possible neural representations of the SIFI that has been addressed in numerous studies. Early electroencephalography (EEG) measurements had shown that activation in primary visual cortex is correlated with the perception of the illusion (e.g., Shams, Kamitani, Thompson, & Shimojo, 2001; Mishra, Martinez, Sejnowski, & Hillyard, 2007). In a recent review summarizing results from EEG, MEG (magnetoencephalography), and TMS (transcranialmagnetic stimulation) studies, Keil (2020) concludes that both early and late processing stages, at different hierarchies in subcortical and cortical brain regions, are involved in producing the SIFI. Most relevant for the TWIN–SIFI model are studies finding a relationship between neural oscillations, in particular individual alpha frequency (IAF) between 8 and 14 Hz (i.e., 120–60 ms cycle), and the probability of an illusion: for example, Keil and Senkowski (2017) showed that the length of the individual alpha band cycle in occipital cortex provides a “window of opportunity” for audiovisual integration such that the length of this window is positively correlated (across participants) with a higher probability of the illusion, which is in correspondence with the TWIN-SIFI model.

An interesting hypothesis that may even suggest a possible extension of TWIN-SIFI comes from studies in Cecere, Rees, and Romei (2015). Applying mild transcranial alternating current (tACS) at and around each individual’s alpha frequency, they confirmed the relation between alpha oscillations and the temporal window of the SIFI. Specifically, the authors then hypothesize that perception of the second flash is the result of an increase in cortical excitability triggered in visual cortex by the second beep, if the beep is presented within an alpha cycle of the flash. In a subsequent study, Stiles, Tanguay Jr., and Shimojo (2020) propose that the likelihood that the illusion actually occurs should depend on (i) how strong the initial visually-driven activation is and (ii) how strong the multisensory activation of visual regions is from auditory brain regions. An extension of TWIN-SIFI would be to add a random variable, *V* ^*t*^, say, representing the illusory flash triggered by the second beep, to each of the 3 events *E*_*i*_(*τ, ω*) (*i* = 1, 2, 3). For example, *E*_1_(*τ, ω*) would become

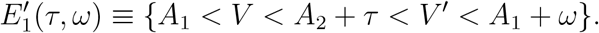

In this case, the visual cortical activation would be strong enough to realize within the temporal window. The typically observed variability in the illusion to occur would be due to variability in the strength of the visual cortical activation from the first real flash.

## Acknowledgments

This work was partly supported by DFG (German Science Foundation) grants to H. Colonius (CO 94/8-1) and A. Diederich (DI 506/18-1).

## Appendices

### Appendix A: Exponential model with unequal *λ* parameters

Relaxing the assumption of equal intensity parameters (*λ*) in the exponential model for events *E*_*i*_, (*i* = 1, 2, 3), the probability of integration with *ω >* 0 becomes (using Mathematica 13.3©):

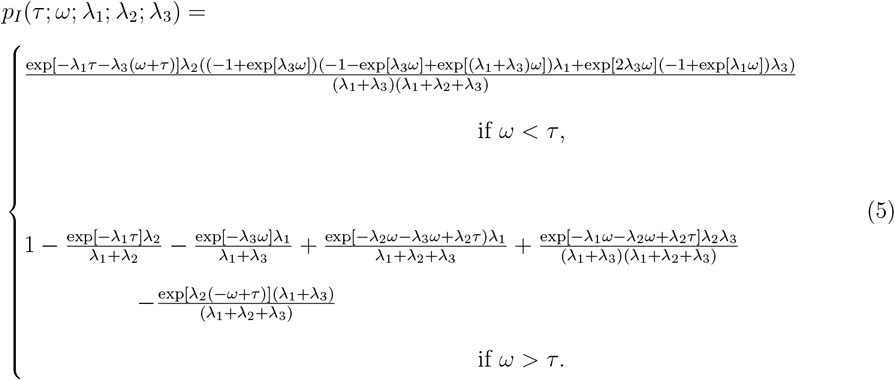

## Appendix B: Parameter estimation and exponential model fits for hearing-aid experiment (Gieseler et al., 2018)

There were 10 data points (illusion/no illusion) per subject per condition. Illusion frequencies were merged across all participants, separately for each SOA condition and groups HU and NU. Table 1 lists the parameter estimates from minimizing the weighted sum of squares between predicted and observed frequencies using Matlab ^©^routine fminsearchbnd.

**Table 1.** Parameter estimates and fit indices for 5 different versions of the exponential model (see text for details).

| Model | $\lambda$ | $\omega$ | $\beta$ | $\lambda$ | $\omega$ | $\beta$ | obf ($\chi^2$) | df | RMSEA |
| --- | --- | --- | --- | --- | --- | --- | --- | --- | --- |
| λ-ω-β | 0.009 | 174 | 0.139 | | | | 214 | 13 | 1.02 |
| $\lambda_{HU/NU}$-ω-β | 0.010 | 183 | 0.136 | 0.006 | | | 166 | 12 | 0.93 |
| λ-$\omega_{HU/NU}$-β | 0.009 | 218 | 0.119 | | 145 | | 96 | 12 | 0.68 |
| λ-ω-$\beta_{HU/NU}$ | 0.009 | 173 | 0.213 | | | 0.048 | 30 | 12 | 0.31 |
| λ-$\omega_{HU/NU}$-$\beta_{HU/NU}$ | 0.009 | 190 | 0.191 | | 163 | 0.057 | 17 | 11 | 0.19 |

Five models (column 1) were considered that varied with respect to the number of parameters: for example, Model *λ*-*ω*-*β*_*HU/NU*_ refers to a model featuring *λ* and *ω* parameters that are identical for groups HU and NU, but separate *β* parameters for HU and NU. The last column lists the root mean square error of approximation (RMSEA) value for each model defined as

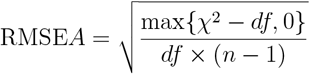

with *n* = 16 the number of observations. The best-fitting model (last row) has 5 parameters and both a larger window of integration and a higher bias value for the HU group compared to the NU group.

## Footnotes

1 When too small SOA (*τ*) values are selected in an experiment, *P* (*A*_1_ *< A*_2_ + *τ*) = 1 may no longer hold. Note, however that this implies that events *{A*_2_ + *τ < A*_1_ *< V < A*_2_ + *τ* + *ω}* and *{A*_2_ + *τ < V < A*_1_ *< A*_2_ + *τ* + *ω}*, may also generate integration and would make derivations more complex but feasible.

2 A non-linear alternative to introduce bias is to assume *p*(*τ* ; *ω, β*) = *p*_*I*_ (*τ* ; *ω*)^1−*β*^ ; however, limit behavior towards 0 and 1 would be the same.

3 Only small SOA values (about 70–120) combined with large window widths (about 300–370) slightly increase integration probability (see upper left corner of Figure 4). The white line on the surface corresponds to the case of *τ* = *ω* that can be neglected here.

4 The distinct role of mild hearing loss on audiovisual integration and the significance of these changes for speech intelligibility have been investigated at the Oldenburg lab for some time (Diederich, Colonius, & Schomburg, 2008; Schulte et al., 2020; Rosemann, Gieseler, Tahden, Colonius, & Thiel, 2021).

5 Stiles et al. (2018) report a second illusion, “invisible audiovisual rabbit”, where a beep-flash pair following a real flash suppresses the perception of the earlier flash.

